# Non-canonical EGFR signaling is essential for MAPK-mediated apical extrusion of epithelial cells

**DOI:** 10.1101/2025.05.16.654585

**Authors:** Paola Molina, Mikiyas Daniel, Jason Wang, Ian Macara

## Abstract

Individual epithelial cells that acutely express oncogenes such as Ras or Src are extruded apically from monolayers of wildtype cells. Multiple signaling networks have been implicated but the extrusion mechanism is still not fully understood. We examined extrusion of mammary epithelial cells caused by acute induction of oncogenic Ras(Q61L). As reported by others, Ras-dependent extrusion requires downstream activation of ERK but not AKT. Unexpectedly, however, extrusion was completely blocked by Erlotinib, which inhibits Epidermal growth factor receptor (EGFR) activity, or by deletion of the receptor. In pancreatic and lung cancers, EGFR is required for full activation of Ras and consequent ERK activation. However, inhibition or deletion of EGFR had no impact in our system on Ras(Q61L)-GTP levels or ERK phosphorylation. Importantly, receptor function was cell autonomous, because EGFR expression was not required in surrounding WT cells but was essential in the Ras(Q61L) cells, yet did not act through the canonical ERK signaling pathway. Deletion of Ras exchange factors Sos1/2 did not block cell extrusion. Moreover, expression of a constitutively active MEK mutant, instead of Ras, was sufficient to drive extrusion, and EGFR inhibition or knockout in these cells blocked extrusion, with no change in phospho-ERK levels. Notably, acute expression of Ras triggered internalization of E-cadherin, which was partially blocked by inhibition of EGFR. Knockout of E-cadherin was alone sufficient to promote extrusion. Together, these data demonstrate an unanticipated requirement for noncanonical EGFR signaling in cancer cell extrusion, which may act in part through the promotion of E-cadherin endocytosis.

**Statement of Significance:** Apical extrusion of cells acutely expressing oncogenic Ras requires EGFR activity through a noncanonical pathway, independent of ERK and AKT signaling, that promotes E-cadherin internalization from adherens junctions.

## INTRODUCTION

Organismal development and health depend on the presence of surveillance systems that can recognize and respond to aberrant cells^1^. Most animal tissues are epithelial, and epithelia are the most common source of human cancers. A variety of epithelial-intrinsic error correction systems have evolved to remove aberrant cells, including cell-cell competition and interface surveillance^1–3^, which depend on the detection of differences between cell neighbors mediated by intercellular contacts. One important example of a response to aberrant cells is the extrusion of individual cancer cells from wild type epithelial cell monolayers or tissues^4, 5^. This process depends on interface surveillance as demonstrated by the fact that extrusion does not occur if all the cells in the monolayer are transformed, but only if a transformed cell is surrounded by wild type cells. Ras-dependent apical extrusion involves multiple changes in both the transformed cell and its nearest neighbors. These include changes in actin dynamics and actomyosin contractility as well as in metabolism, Ca^2+^ signaling, reactive oxygen production, and EphA2/Ephrin-triggered cell repulsion^3, 4, 6–8^; however, the complete mechanism of extrusion remains elusive. One aspect of Ras-driven extrusion that has not previously been considered is a role for Epidermal growth factor receptor (EGFR) signaling.

Ras mutations are frequent in lung and pancreatic cancers but in several mouse models of these cancers, oncogenic Ras activity requires EGFR activation and tumor growth is reduced by EGFR ablation^9–12^. Using a lung cancer cell line, A549, which expresses an oncogenic K-Ras mutant, knockout of EGFR reduced the level of activated (GTP-bound) Ras^10^. EGFR was also shown to be required for tumorigenesis in a KRas(G12D) pancreatic cancer model, and ablation of EGFR reduced downstream Ras-GTP and ERK phosphorylation levels by ∼2-fold^9^. Moreover, in patient-derived organoids of KRas-driven colorectal cancers, EGFR activity was essential to promote ERK phosphorylation and tumor cell proliferation^12^. EGFR was also found to be needed for the growth and survival of Ras-initiated squamous cell carcinoma^13, 14^ and melanomas^15^. The SOS1/2 guanine nucleotide exchange factors, which activate WT Ras, function downstream of the EGFR and other receptor tyrosine kinases but also independently of EGFR through allosteric interaction of SOS with oncogenic K-Ras, which contributes to pancreatic cancer cell growth^16^. Thus, while EGFR might in principle be required for tumorigenesis in the above examples, by promoting activation of the WT Ras alleles in the tumor cells, oncogenic Ras would seem capable of fulfilling this role and obviate any necessity for EGFR signaling. Moreover, oncogenic Ras can desensitize signaling from the EGFR^17^.

Based on these data, it seemed unclear whether Ras-driven cell extrusion would require EGFR activity or not, even though the necessity for EGFR signaling in KRas tumor models is incontestable. We used murine mammary epithelial cells (Eph4) as a model because these cells form highly polarized monolayers in culture and are easily manipulated using Cas9-mediated gene editing. We first confirmed that in this model the acute expression of oncogenic Ras(Q61L) triggers apical extrusion and that extrusion requires MEK activity but not PI3K activity. Notably, inhibition of EGFR efficiently blocked extrusion. Ablation of EGFR in the Ras-expressing cells but not WT neighboring cells also blocked extrusion. Surprisingly, however, inhibition of EGFR had no significant effects on Ras-GTP or phospho-ERK levels. Moreover, deletion of the Ras exchange factors SOS1/2, which act downstream of EGFR in the canonical signaling pathway, also had no effect on extrusion. Induced expression of a constitutively active MEK was as effective as oncogenic Ras in triggering extrusion and this response was blocked by inhibition or loss of EGFR, strongly suggesting that EGFR plays a role independent of the SOS->Ras->Raf->MEK->ERK signaling pathway. Notably, expression of Ras(Q61L) induced the internalization of E-cadherin, which was partially blocked by the inhibition of EGFR.

## RESULTS

### Ras-dependent apical extrusion of epithelial cells is dependent on EGFR and MEK activity

We first demonstrated that we could recapitulate the known ability of oncogenic Ras to trigger cell extrusion^4^, using the Eph4 mammary epithelial cell line and a lentiviral, Dox-inducible HRas(Q61L) fused at its N-terminus to eGFP via the self-cleaving peptide P2A (Fig 1 A). When these cells were co-cultured at a 1:50 ratio with WT Eph4 cells that were marked with mApple, addition of Dox (100 μg/ml) induced apical extrusion within 15 hrs (Fig 1 B). About 75% of the Ras cells were extruded by 24 hrs Dox treatment (Fig 1 C). No extrusion was detected for cells expressing eGFP alone (EV, empty vector, Fig 1 A – C). As expected, induction of oncogenic Ras activated the ERK pathway, increasing phospho-ERK levels (Fig 1 D, E). Maximal phosphorylation occurred after ∼15 hrs Dox treatment, even though Ras levels continue to rise through 24 hrs (Fig 1 A).

**Figure 1.**
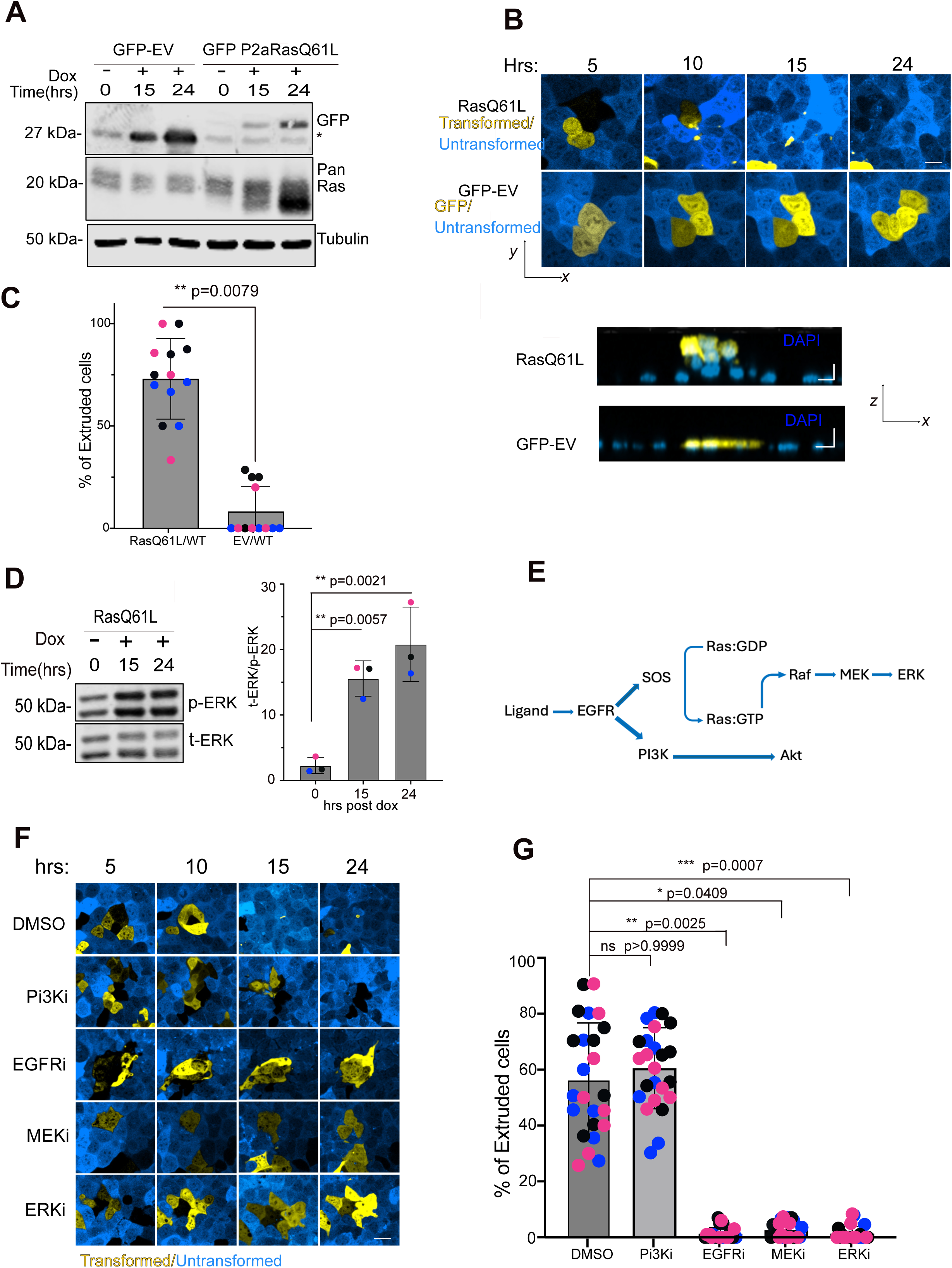
MAPK and EGFR activity are required for epithelial cell extrusion. **(A)** Representative immunoblot for GFP (Rabbit) and Pan-Ras(mouse) showing that GFP is expressed in both cell lines, but Ras expression increased only in RasQ61L cell line after Dox induction. Cells were harvested 0, 15 and 24 hrs after Dox addition. **(B)** Confocal images of the basal plane and orthogonal view of Eph4 murine mammary epithelial monolayers containing cells with inducible GFP-P2a-RASQ61L or GFP, mixed with mApple-tagged cells and cultured at a 1:50 ratio. Cells were treated with doxycycline (Dox, 100μg/mL) to induce the GFP constructs, and imaged over 24 hrs. Scale bar = 10μm. **(C)** Quantification of GFP+ cells apically extruded from the monolayer (N=3 independent experiments). Mean +/- SD. Mann-Whitney test was applied (two-tailed). **(D)** Representative immunoblot for phospho- and total ERK showing increased activity in RasQ61L cells after Dox addition. Quantification of ERK phosphorylation over total ERK for N=3 independent experiments. (Mean +/- SD). One-way ANOVA test was applied. **(E)** Simplified schematic of the EGFR signaling pathway. **(F)** After Dox induction, confocal images of mutant Ras cells/normal cells mixtures were obtained after 5 - 24 hrs + Dox in the presence of DMSO, Pi3K inhibitor LY20049 (10μM), EGFR inhibitor Erlotinib (10 μM), MEK inhibitor U0126 (10 μM) and ERK inhibitor SCH772984 (10 μM). Scale bar=10μm. **(G)** Quantification of extruded GFP+ RasQ61L cells. In the presence of MEKi, ERKi and EGFRi, Ras cell extrusion was significantly reduced in all three conditions (N= 3 independent experiments. Data +/- SD). Kruskal-Wallis test was performed.

We next tested the dependency of apical extrusion on various signaling pathways. Inhibition of the PI3K->AKT pathway using LY294002 had no effect on extrusion (Fig 1E-G; Supplementary Fig S1 A) as has been reported previously^4^, but inhibition of the MAPK pathway using either the MEK inhibitor U0126 or ERK inhibitor SCH772984 suppressed ERK phosphorylation (Supplementary Fig S 1 B) and completely blocked extrusion (Fig 1 F, G). Notably, inhibition of the EGF receptor with Erlotinib also completely suppressed extrusion (Fig 1 F, G, Supplementary Fig S1 C). This result was surprising, but several papers have reported that in mouse tumor models, oncogenic Ras activity (GTP-binding) and ERK phosphorylation are significantly reduced by ablation or inhibition of the EGFR, paralleled by reduced tumorigenesis^9–15, 18^. In some cases^9^, but not in all, this reduction might be ascribed to reduced GTP-binding by the WT allele retained in the KRas(G12D) model.

### Inhibition of or deletion of EGFR does not impact Ras-GRTP levels or ERK phosphorylation

To investigate whether EGFR inhibition causes any decrease in Ras signaling, we first measured Ras-GTP levels using a GST fusion of Raf-RBD. As shown in Fig 2 A, B, Ras-GTP levels were increased by induction of Ras(Q61L) but were not reduced by subsequent addition of Erlotinib. Moreover, addition of Erlotinib had no effect on the increased phosphorylation of ERK caused by Ras(Q61L) induction (Fig 2 C, D). Note that significant ERK phosphorylation is detected in WT cells in the absence of inhibitor, which is suppressed by Erlotinib treatment (Fig 2 C, D).

**Figure 2.**
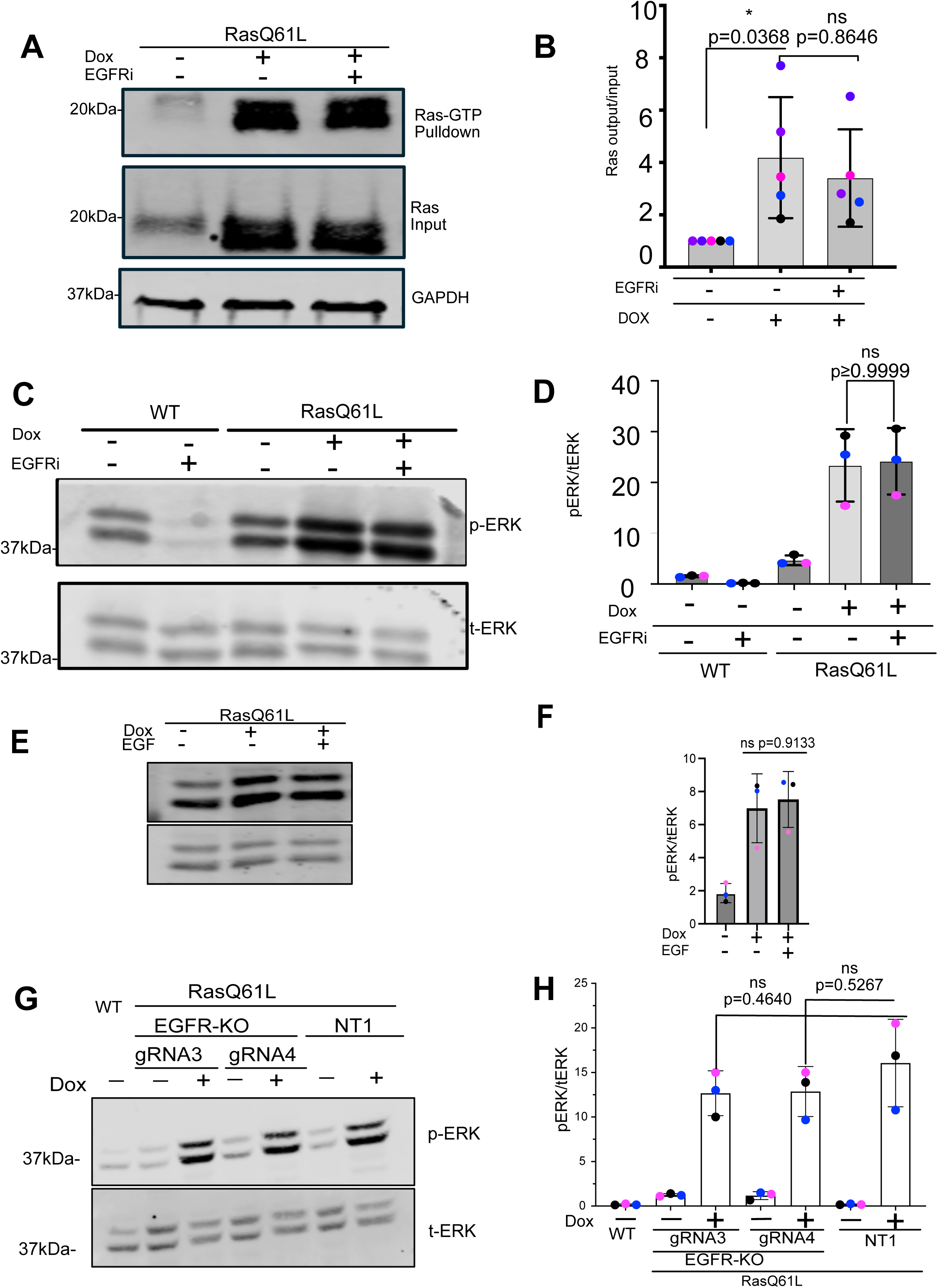
EGFR does not increase Ras GTP-loading or ERK phosphorylation foll owing induction of HRas(Q61L). **(A)** Ras-GTP loading assay. HRas(Q61L) was induced by +Dox for 24 hrs +/- Erlotinib to block the EGFR. Ras-GTP was captured on Raf-RBD-GST beads from cell lysates and analyzed by immunoblot for pan-Ras. Total Ras was analyzed from 5% of the lysates, with GAPDH as a loading control. **(B)** Quantification of the RasGTP/Ras input ratio (N= 5 biological replicates. Data +/- SD). One-way ANOVA with Sidak’s multiple comparison test. **(C)** total ERK (t-ERK) and phospho-ERK (p-ERK) immunoblot from the same gel on cell lysates from starved WT Eph4 cells or cells induced by + Dox for 24 hrs to express HRas(Q61L), +/- 10 μM Erlotinib or vehicle. **(D)** Quantification of the blots demonstrates that inhibition of EGFR activity reduces p-ERK levels in the WT control cells but has no effect on p-ERK in cells expressing HRas(Q61L). (N= 3 independent experiments. Data +/- SD). Mann-Whitney test was performed to compare HRas(Q61L) =/-ERLOT. **(E)** Cells lacking EGFR show the same increase in p-ERK in response to oncogenic Ras expression as control cells (NT1). Cell lysates prepared and blotted at 24 hrs after Dox or vehicle addition. **(F)** Quantification of blot shows that p-ERK levels are increased further by the addition of EGF when oncogenic Ras is activated. (N=3 independent experiments. Data =/-SD. One-way ANOVA test was performed. **(G)** Cells lacking EGFR show the same increase in p-ERK in response to oncogenic Ras expression as control cells (NT1). Cell lysates prepared and blotted at 24 hrs after Dox or vehicle addition. **(H)** Quantification of p-ERK/t-ERK ratios from blots as performed in (E). N= 3 biological replicates. One-way ANOVA, bars show mean +/- SD.

As these experiments were performed in medium lacking serum or added growth factors, they indicate that WT Eph4 cells constitutively release an EGFR ligand that contributes most of the background ERK activation. Importantly, addition of exogenous EGF did not further stimulate ERK phosphorylation above that caused by Ras induction, suggesting that Ras(Q61L) fully activates the MEK/ERK pathway (Fig 2 E, F). Moreover, addition of serum (which does not contain EGF) to the cell culture medium strongly activated ERK phosphorylation independently of the EGFR, as it was not reduced by Erlotinib treatment (Supp Fig S2 A). Nonetheless, Ras-dependent extrusion was still blocked by the inhibitor (Supp Fig S2 B). Taken together, these data suggest that cell extrusion requires both MAPK signaling and a MAPK signaling-independent function of the EGFR.

To more definitively validate these results, we used Cas9-mediated gene editing to delete EGFR in the Eph4-Ras cells, using 2 independent gRNAs, or a non-targeting gRNA as a negative control. Knockout was very efficient (Supp Fig S3 A). However, induction of Ras(Q61L) expression in these cells stimulated ERK phosphorylation to the same level in the presence or absence of the EGFR (Fig 2 G, H). These data suggest that in this acute Ras-induction model, EGFR is not necessary for full activation of Ras or for full stimulation of the MAPK pathway, even though it is essential for apical extrusion.

### EGFR-dependent extrusion is induced by a constitutively active MEKK

The data described above (Fig 1 F, G) demonstrate a requirement for ERK phosphorylation downstream of Ras (Q61L) expression but do not address sufficiency, nor whether other pathways downstream of Ras are important. Therefore, we generated a cell line containing a Dox-inducible, constitutively active MEK mutant (MEKDD), fused to a self-cleaving GFP-P2A as an expression marker (Fig 3 A, B). First, we showed that, as expected, Induction of MEKDD expression had no effect on the PI3K-> AKT pathway, as opposed to Ras(Q61L) expression, which strongly promoted AKT phosphorylation (Fig 3 C, D). Importantly, however, MEKDD expression did trigger ERK phosphorylation, and to a similar degree as did Ras(Q61L) (Fig 3 E, F). Moreover, MEKK expression triggered cell extrusion to a similar extent as oncogenic Ras (Fig 3 G, H). Strikingly, Erlotinib treatment completely blocked MEKDD-driven apical extrusion (Fig 3 I, J). Together, these data conclusively demonstrate that EGFR activity is essential for cell extrusion driven by ERK signaling but through a noncanonical pathway that acts downstream of Ras and does not enhance Ras-GTP or ERK signaling.

**Figure 3.**
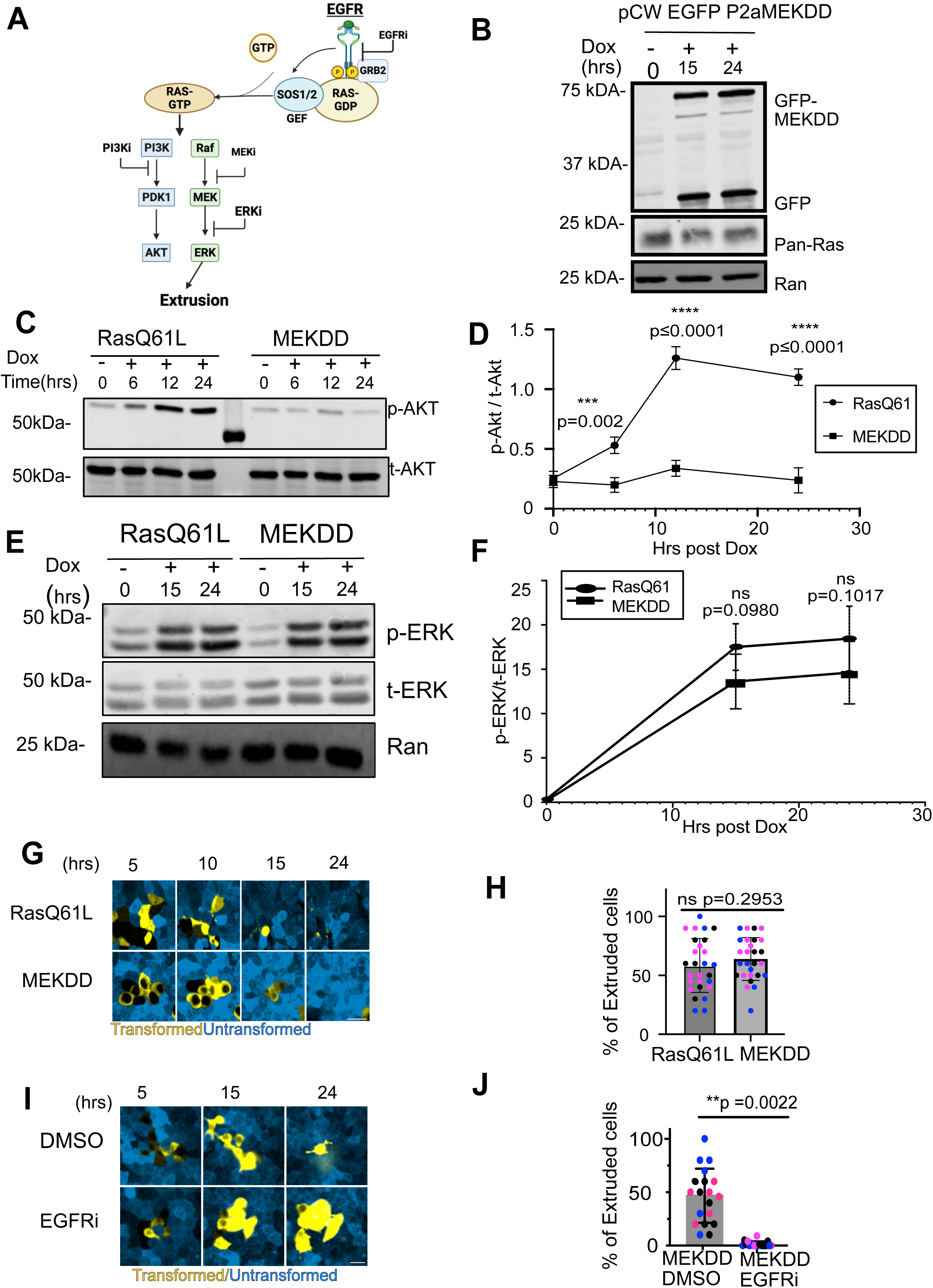
Constitutively active MEKK triggers cell extrusion, which is blocked by EGFR inhibition. **(A)** Graphic summarizing prior experiments using pharmacological inhibitors targeting EGFR (EGFRi), PI3K (PI3Ki), MEK (MEKi), and ERK (ERKi) to assess their roles in cell extrusion. Inhibition of EGFR, MEK, or ERK impaired cell extrusion, whereas PI3K inhibition had no significant effect, highlighting the critical role of MAPK signaling in this process **(B)** Western blot validating the inducible P2A-MEKDD construct. Cells expressing P2A-MEKDD (a constitutively active mutant of MEK) in Dox-inducible GFP vector show GFP expression upon Dox treatment, confirming induction. Ras protein levels remain unchanged, indicating that expression of the MEKDD construct does not affect endogenous Ras levels. **(C)** Western blot analysis of p-AKT and total AKT (t-AKT) levels in cells expressing Dox inducible RasQ61L or MEKDD constructs. Following Dox addition, RasQ61L expression increases p-AKT levels over time, whereas MEKDD expression does not significantly alter p-AKT levels. Total AKT levels remain constant across all conditions. **(D)** Quantification of p-AKT levels in cells expressing Dox-inducible RasQ61L or MEKDD constructs. p-AKT intensities were normalized to total AKT. RasQ61L expression led to a time- dependent increase in p-AKT levels, whereas MEKDD expression had minimal effect. Two-way ANOVA was performed, Data =/-SD(N=3). **(E)** Dox-induced expression of MEKDD promotes phosphorylation of ERK to a similar level as oncogenic Ras. Cell lysates were blotted for p-ERK and t-ERK on the same gel, with anti-Ran as a loading control. **(F)** Quantification of blots performed, N=3 biological replicates, bars show mean +/- 1 SD, compared by 2way ANOVA. **(G)** MEKDD drives extrusion of Eph4 epithelial cells from a monolayer after induction with Dox, over a similar time course to oncogenic Ras. Scale bar= 10 μm. **(H)** Quantification of extrusion at 24hrs post-induction is not significantly different for MEKDD versus Ras(Q61L). Bars show mean +/- 1 SD (N=3); dots of the same color are data for different fields from the same replicate. Mann-Whitney test was performed to compare HRas(Q61L) to MEKDD. **(I)** Inhibition of EGFR with Erlotinib blocks MEKDD induced cell extrusion. Scale bar = 20μm**. (J)** Quantification of data from MEKDD induced cell extrusion +/- Erlotinib. Bars show mean +/- 1 SD (N=3); dots of the same color are data for different fields from the same replicate. Mann-Whitney test was performed to compare MEKDD +/- ERLOT.

### Extrusion requires EGFR in the p-ERK positive cells, not in the surrounding WT cells

Small molecule inhibitors do not distinguish between the two cell types present in the extrusion assay. Therefore, we took advantage of our EGFR knockout cell lines to test if the receptor is needed in the WT cells or the Ras-expressing cells, or both. We expected that receptor would be required in the WT cells based on prior data for MCF10A cells^19^, in which release of the EGFR ligand AREG by oncogenic Raf(V600E) activates ERK in neighboring WT cells^19^. However, to the contrary, we found that extrusion of Eph4 epithelial cells induced for Ras(Q61L) expression was unchanged when they were surrounded by either WT (non-targeting sgRNA, NT1) or EGFR-/- cells transduced with either of 2 independent sgRNAs (Fig 4 A, B, E). We next induced Ras(Q61L) expression in cells deleted for EGFR and surrounded by WT (NT1) cells. In this case, extrusion was completely suppressed (Fig 4 A, C, F). Strikingly, extrusion of cells expressing the active MEKDD was also blocked by EGFR knockout (Fig 4 A, D, G). We conclude, therefore, that EGFR signaling is needed in the extruding cells, not in neighboring cells, and that its function is entirely independent of any effects on Ras-GTP loading or ERK phosphorylation.

**Figure 4.**
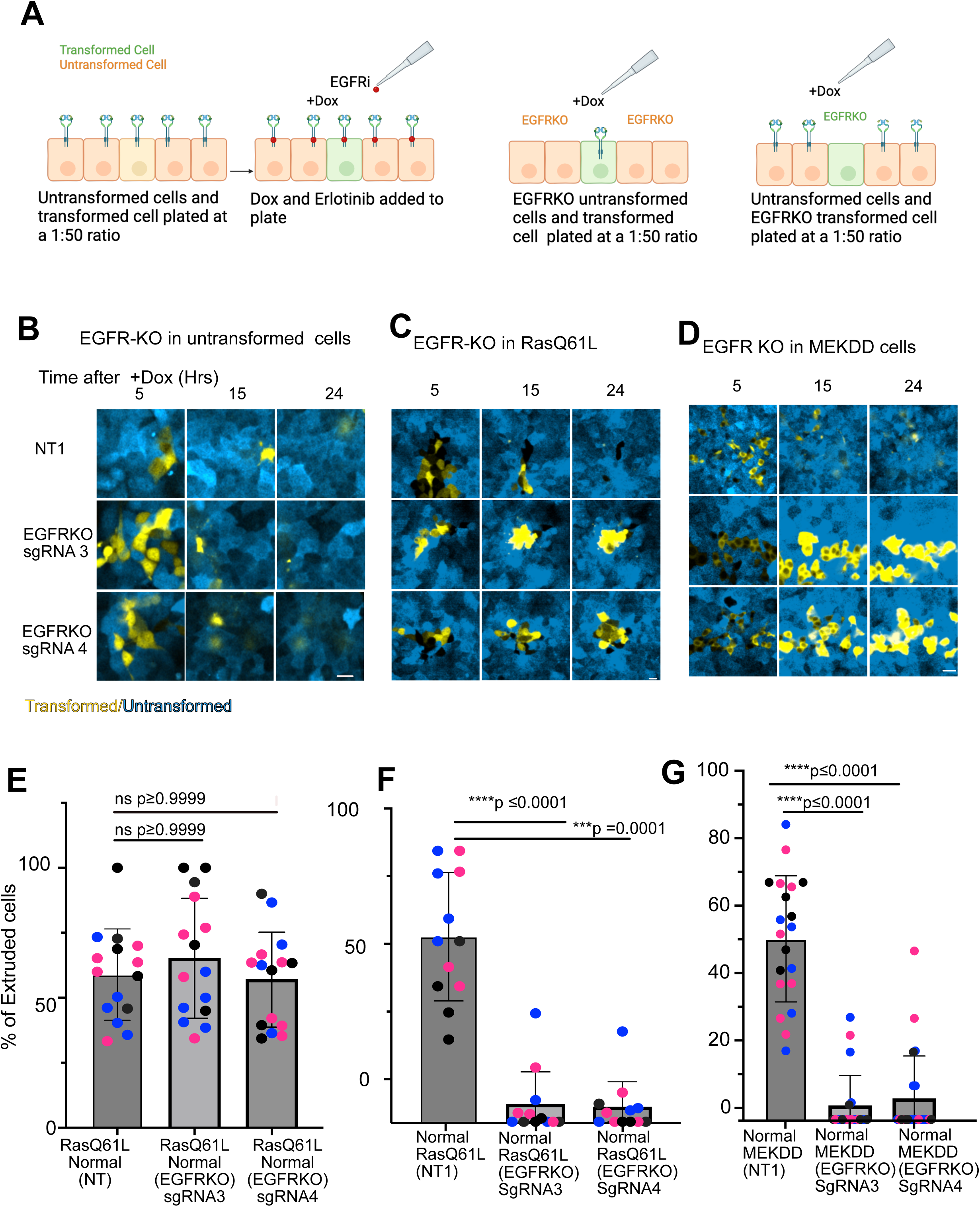
EGFR is required in the Ras(Q61L) and MEKDD cells for extrusion, but not in neighboring cells. **(A)**Schematic of the experimental design to assess the role of EGFR in normal versus transformed cells during extrusion. EGFR was selectively ablated using CRISPR-Cas9 in either the normal or transformed cell population to determine which cell type–associated EGFR activity is required for extrusion. **(B)** Ras(Q61L) cells (yellow) were plated at 1:50 with control cells (blue) deleted for EGFR (EGFR-KO) or a control sgRNA (NT1) and treated with Dox to induce Ras expression. Images were captured at intervals to assess extrusion of the Ras cells. Scale bar = 20 µm **(C)** Ras(Q61L) cells deleted for EGFR (yellow) were plated at 1:50 with WT cells (blue) and treated as in (B). Scale bar= 10 µm. **(D)** MEKDD cells deleted for EGFR were plated as in (C) with WT cells and induced by addition of Dox to induce MEKDD expression. Scale bar=20 µm **(E-G)** Quantification of cell extrusion corresponding to the experimental conditions shown in panels B–D, respectively. Individual data points represent biological replicates (different colors), with multiple points of the same color corresponding to different fields of view. Bars show mean +/- 1 SD (n = 3). Kruskal-Wallis test.

### Knockout of Ras exchange factors SOS1/2 has no effect on extrusion

Ligand-mediated activation of the EGFR causes tyrosine phosphorylation of the C-terminal region of the receptor and the assembly of a signaling hub that activates guanine nucleotide exchange factors for Ras, called SOS1 and SOS220. Based on our previous data, we would predict that deletion of these factors would not impact extrusion. However, earlier studies had demonstrated that SOS2 is required for KRas-dependent transformation by promoting activation of endogenous WT HRas^21^. The underlying mechanism depends on an allosteric site on SOS for KRas that stimulates its guanine nucleotide exchange activity towards the WT H- (and N-) Ras^16^. Therefore, it seemed possible that SOS1/2 might indirectly contribute to cell extrusion either by activating endogenous Ras isoforms or through binding to other components of the signaling hub on EGFR.

We knocked out each of the two isoforms of SOS in the inducible KRas(Q61L) cell line and isolated clones negative for both (Fig 5 A). Although addition of EGF to the WT (NT1,2) cells triggered the expected increase in EGFR tyrosine phosphorylation and p-ERK, the change in p-ERK was much reduced in the knockout clones (Fig 5 A, B). However, after induction of oncogenic KRas expression by Dox addition, extrusion proceeded as for the NT 1,2 cells over the course of 24 hrs (Fig 5 C, D). We conclude that even though EGFR kinase activity is required, downstream signaling through the Grb2/SOS signaling hub is dispensable for cell extrusion.

**Figure 5.**
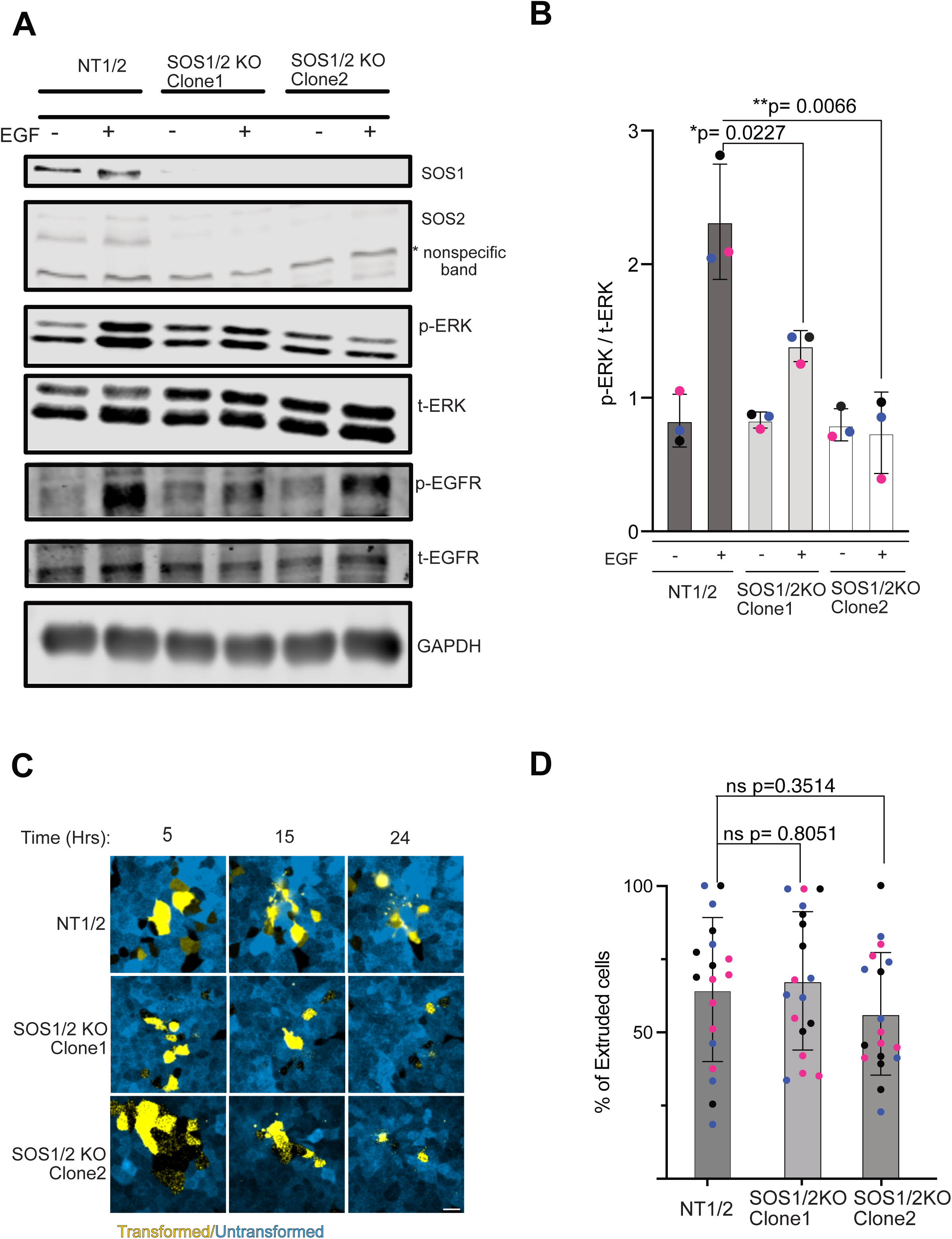
Deletion of Ras exchange factors SOS1 and SOS2 does not block Ras(Q61L)-dependent extrusion. **(A)**SOS1 and SOS2 were deleted from the Ras(Q61L) cell line by Cas9-mediated gene editing. Cells were selected using puromycin and blastocidin, then cloned to obtain cell lines lacking both exchange factors. Two independent clones were blotted for SOS1 and SOS2 expression, and for p-ERK levels after EGF addition (in the absence of Dox). They were also probed for tyrosine phosphorylation of the EGFR. GAPDH was a loading control. **(B)** Quantification of p- ERK levels in SOS1/2-KO and control cells with or without EGF stimulation. Groups were compared using an unpaired two-tailed t-test. Bars represent mean +/- 1 SD (n = 3). **(C)** Cell extrusion assay showing fields of Ras(Q61L) cells (yellow) either expressing a non-targeting sgRNA or deleted for SOS1+SOS2 and imaged at intervals after treatment with Dox to induce Ras expression. Scale bar= 20µm **(D)** Quantification of cell extrusion efficiency in SOS1/2 knockout clones compared to the NT1 control line. Bars represent mean +/- 1 SD (n = 3). Groups were compared using a two-tailed unpaired t-test.

### E-cadherin is rapidly internalized after induction of Ras(Q61L), but internalization is blocked by EGFR inhibition

Staining for E-cadherin after induction of oncogenic Ras in the Eph4 cells revealed a substantial internalization of the adherens junction protein between 10 -24 hrs after Dox addition (Supp Fig S4 A-C). During this period there was no reduction in E-cadherin levels (Fig S4 D).

Internalization was completely blocked by treatment of the cells with MEK inhibitor (U0126), consistent with the process being driven by ERK phosphorylation (Fig S2 D - F). To test whether EGFR is required for E-cadherin internalization, we used Eph4 cells expressing E-cadherin-RFP and created timelapse movies of the cells treated +/- Erlotinib prior to induction of oncogenic Ras. The effect on E-cadherin-RFP association with adherens junctions was quantified by drawing a rectangular grid over each frame of the videos and computing the frequency with which the grid lines intersect with RFP fluorescence above a background threshold (the intersection parameter) (Supp Fig S4 G). Removal of E-cadherin-RFP from the junctions by endocytosis increases the distance between intersections. We noticed that inhibition of EGFR maintained the appearance of the E-cadherin at junctions (Fig 6A), and this was validated by the intersection parameter, which was significantly (though not completely) reduced by Erlotinib (Fig 6 B). Thus, blocking EGFR activity partially suppresses E-cadherin internalization.

**Figure 6.**
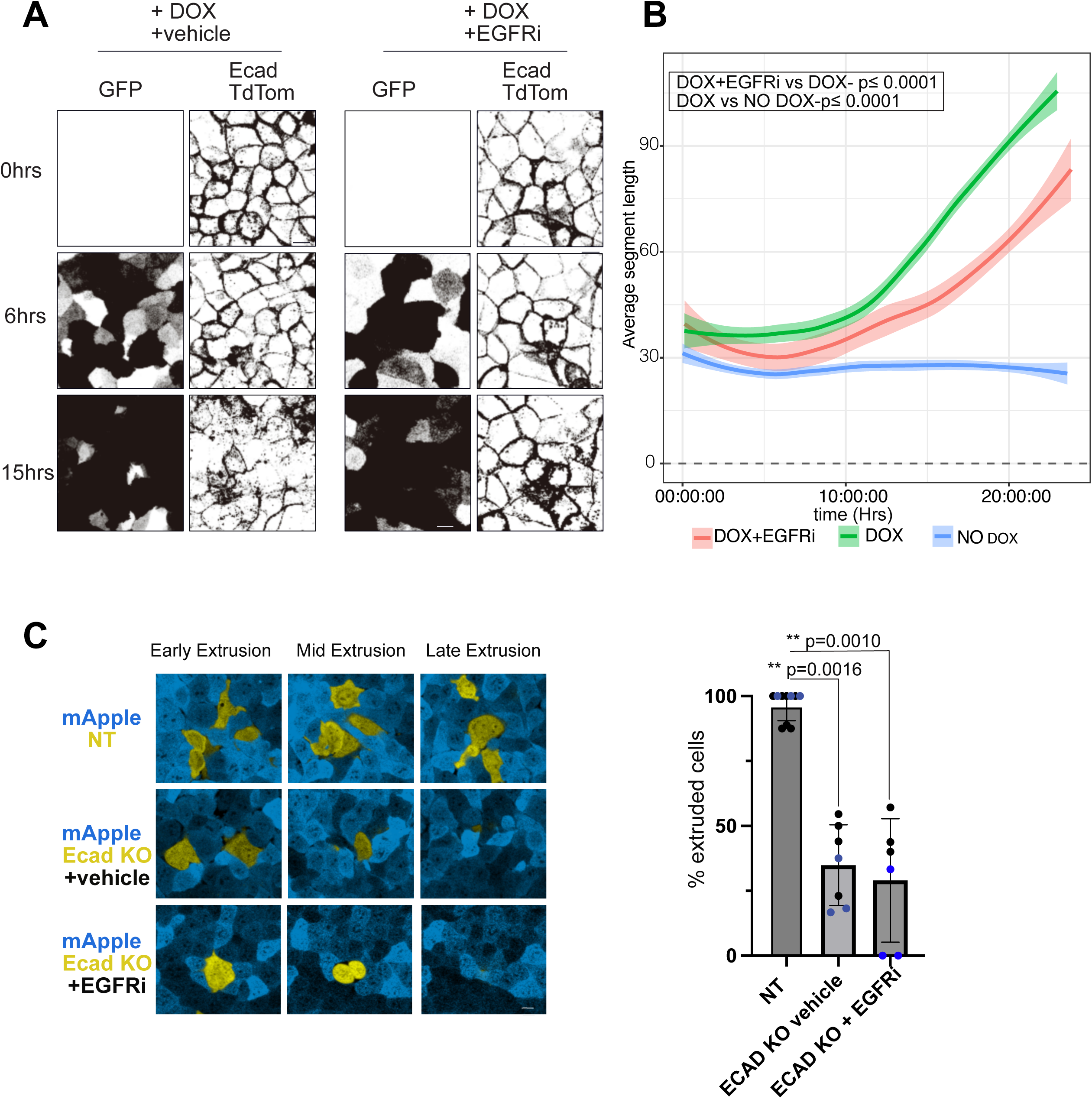
E-cadherin is internalized by Ras activation through an EGFR-dependent mechanism. **(A)** Eph4 cells expressing E-cadherin-tdTom and Dox-inducible Ras(Q61L), with or without erlotinib treatment, were imaged for up to 24 hrs after Dox addition. MaxIP images show that cells treated with erlotinib exhibited less E-cadherin disruption compared to untreated cells. Scale bar = 10 µm. **(B)** Analysis of segment length between E-cadherin junctions over time, in which longer average segment lengths indicate greater junctional disruption (see Supp Fig S4 F). Lowess curves represent each condition (n=3). A repeated-measures ANOVA revealed a significant difference from control in junction integrity between treatments (****p ≤ 0.00001). **(C)** Confocal imaging of a heterogeneous mixture of E-cadherin knockout (KO) and wild-type cells over 24 hrs. Both E-cadherin KO and non-targeting (NT) control cells carried a Halotag vector and were stained with JF646 dye. E-cadherin KO cells were preferentially eliminated from the monolayer, while NT cells remained. Scale bar = 10 µm. Extrusion was quantified from videos (N=2 biological replicates, 3 fields of view per replicate). Bars represent mean +/- 1 SD (n = 3). Groups were compared using a two-tailed unpaired t-test.

Does loss of E-cadherin impact extrusion? To test this hypothesis, we knocked out E-cadherin from Eph4 cells and mixed the cells with WT cells at a 1:50 ratio, then imaged them over time. Strikingly, the E-cadherin null cells rapidly extruded from the monolayer and were lost into the medium (Fig 6 C). Thus, we conclude that loss of E-cadherin from adherens junctions is sufficient to enable extrusion, in agreement with other studies^22, 23^. Partial retention of E-cadherin at junctions by inhibition of EGFR might, therefore, act to suppress extrusion driven by oncogenic Ras.

## DISCUSSION

The extrusion of cancer cells embedded in wild type epithelial sheets has been studied for several years and has been proposed in some contexts to function as a defense against cancer^24, 25^. It is closely related to other quality control processes by which abnormal or dying cells are eliminated from a tissue, such as cell competition and interface surveillance^8, 26^. Apical extrusion of oncogene-transformed cells from several epithelial tissues has been observed in vivo as well as in cell culture and multiple signaling pathways have been implicated but the fundamental mechanism remains to be fully understood^6, 24, 27^. A classic approach has been to induce expression of an oncogene sparsely within a culture of epithelial cells such as MDCK or MCF10A.

We have explored the apical extrusion of Eph4 murine mammary epithelial cells induced to express Ras(Q61L) by addition of Doxycycline. Consistent with previous studies^4^, we found that Ras-driven extrusion is dependent on ERK phosphorylation but not AKT activation. Strikingly, however, inhibition of the EGF receptor tyrosine kinase activity also completely blocked apical extrusion, even though this receptor functions upstream of Ras. Knockout of EGFR in the Ras cells, but not the WT neighboring cells, also blocked extrusion. This observation suggests a fundamentally different mechanism from one reported for MCF10A cells^19^, in which the ADAM17 protease is shed from Ras-expressing cells to release the EGFR ligand AREG, which then stimulates ERK activation in neighboring cells, triggering cell migration that drives Ras cell extrusion.

Multiple papers have demonstrated that EGFR is required in pancreatic and lung cancer models driven by oncogenic KRas, and that oncogenic Ras activity (GTP-binding) and ERK phosphorylation are substantially reduced by ablation or inhibition of the EGFR, paralleled by reduced tumorigenesis^9–15^. In some cases, but not in all, this reduction might be ascribed to reduced GTP-binding by the WT allele retained in the KRas(G12D) model used in the cancer models. In other cases, activation of other endogenous, WT isoforms of Ras (HRas, NRas) by EGFR has been proposed to further stimulate the ERK pathway^17^. Additionally, GTP-loading of oncogenic Ras might require SOS1/2 nucleotide exchange activity stimulated by EGFR^21^.

However, oncogenic Ras can also desensitize signaling from the EGFR. Notably, in our model system, ablation of the SOS exchange factors had no impact on extrusion efficiency. Interestingly, an analysis of Ras(G12V)-driven tumorigenesis in *Drosophila* also demonstrated that knockdown of Sos had no effect on tumor overgrowth, even though Egfr signaling was essential28.

A key difference between our study and the mouse and human cell line cancer models is that these models all involved chronic KRas expression, while in our system Ras(Q61L) was expressed for only a few hours to initiate cell extrusion, and we found that inhibition of EGFR or ablation of the EGFR gene caused no reduction in Ras(Q61L)-GTP and no reduction of phospho-ERK. Moreover, instead of oncogenic Ras, we were able to induce extrusion simply by expression of a constitutively active MEK mutant, MEKDD, directly upstream of ERK. Taken together, these data argue for a novel, non-canonical function of EGFR activity in the Ras-expressing cells, independent of ERK and AKT signaling pathways, to permit extrusion.

Our data further suggest that EGFR activity is necessary for efficient E-cadherin removal from adherens junctions through endocytosis driven by oncogenic Ras. Deletion of E-cadherin from WT Eph4 cells was sufficient to drive extrusion in the absence of any EGFR signaling, consistent with other studies^22, 23^, but the role of E-cadherin is context specific, as loss of E-cadherin expression in other systems can lead to defects in apoptotic extrusion^29^. In Eph4 cells, ERK activity is required for internalization of E-cadherin, but another, non-canonical EGFR-specific mechanism is also involved. We speculate that a threshold for E-cadherin internalization is needed to drive Ras-dependent extrusion, which is prevented by EGFR inhibition. It will be of interest to probe for the underlying mechanism of this effect.

## ACKNOWLEDGEMENTS

This work was supported by NIH grant GM070902 to IGM. PM was supported in part by training grant T32119925. We thank members of the Macara lab for helpful advice, and especially Christian de Caestecker for the junction integrity algorithm used in Fig 6.

## METHODS

### Cell lines

Mouse mammary EpH4 cells were provided by Dr. Jürgen Knoblich (Institute of Molecular Biotechnology, Vienna, Austria). EpH4 cells were cultured in Dulbecco’s Modified Eagle Medium (DMEM) (ThermoFisher Scientific, Waltham, MA) supplemented with 10% Gibco™ Fetal Bovine Serum (FBS), and incubated at 37°C and 5% CO_2_. 293T cells were cultured in DMEM, 10% FBS, and 100U/mL Penicillin and Streptomycin.

### Plasmid construction and lentivirus production

Stable cell lines expressing RasQ61L and MAPKDD were made by lentivirus transduction. Sequences encoding P2aRasQ61L and P2aMAPKDD were cloned into the doxycycline-inducible pCW-eGFP lentivector (Addgene #162823). P2a sequences were added by PCR to RasQ61L and MAPKDD open reading frames, then ligated into pCW-eGFP downstream of the EGFP using 5’ BsrG1 and 3’ BamH1 restriction sites, to express self-cleaving GFP-P2a-RasQ61L or -MEKDD fusion proteins. The sgRNAs used in this study are listed in the Resources Table. gRNAs were designed using Benchling and ligated into lentiCRISPRv2 (Addgene # 52961) at the BsmBI restriction sites using the Zhang lab protocol^30^. pLVTHM-mApple cells were generated using plasmids described in Ahmed and Macara^31^.

To produce lentivirus, 293T cells were transfected using calcium phosphate with packaging plasmids psPax2 and pMDG2-VSVG along with the desired plasmid to be packaged in the lentiviral genome. Medium was changed after overnight incubation, and virus-containing media was collected after 48hr. Virus was concentrated using Amicon 100k centrifugal filters.

### In vitro Extrusion assay

Eph4 cells were cultured in DMEM, 10% FBS (fetal bovine serum), and 100U/mL Penicillin & Streptomycin. Stable cell lines expressing RasQ61L or MEKDD, or eGFP alone as a control, were mixed at a 1:50 ratio with WT cells expressing mApple, totaling 1.5x10^5 single cells. They were plated onto 8-well chambered coverslips and allowed to settle overnight. The next day they

were treated with 10μg/mL of doxycycline to induce expression of RasQ61L or MEKDD +/1 inhibitors or vehicle at the concentrations listed in the Table. To quantify the percentage of cells that were extruded, dishes were fixed 24 hrs post doxycycline induction and GFP-positive cells above the monolayer (extruded cells) and in the monolayer were counted and calculated as the ratio of (extruded GFP+ cells)/(total GFP+ cells). Extruded cells remained attached to the top of the monolayer and were not lost during the washing/antibody incubation steps, as judged by comparison of the total GFP+ cells (extruded + monolayer) to the eGFP control (monolayer only).

### E-cadherin imaging and E-cadherin KO cell extrusion

To examine the impact of RasQ61L and EGFR inhibition on E-cadherin localization, we generated Eph4 cell lines expressing an E-cadherin-dsRed fusion, plus eGFP alone, or eGFP-P2a-RasQ61L. These cells were plated onto chambered coverglasses, treated with doxycycline, and imaged live for up to 24 hrs by confocal microscopy. Some cultures were also treated with Erlotinib at 10 μM to inhibit EGFR activity. The integrity of the junctions was quantified using a custom Python script to generate an intersection parameter.

Eph4 cells deleted for E-cadherin were created by lentiviral expression of Cas9 plus sgRNAs targeting murine E-cadherin exons (or a non-targeting gRNA as a control). These cells also express a Halotag, labeled with far-red JF646 (Janelia Research Campus).

### The Ras activation assay

The Ras activation assay was performed using the Ras Activation Assay Biochem Kit (Cat. # BK008, Cytoskeleton, Inc.) following the manufacturer’s instructions. Cells were cultured in growth medium until reaching 70% confluency. Serum starvation was performed by replacing the medium with serum-free medium overnight prior to doxycycline and Erlotinib treatment. Cells were treated and incubated for 24 hrs. Cells were then washed twice with ice-cold PBS and lysed using the provided Cell Lysis Buffer supplemented with protease inhibitor cocktail and Phosphostop. Lysates were collected by scraping and clarified by centrifugation at 10,000 x g for 1 min at 4°C. Protein concentration was determined using the Precision Red Advanced Protein Assay Reagent (Cat. # ADV02) following the manufacturer’s protocol. Lysates were incubated with Raf-RBD beads for 1 hr at 4°C on a rotator. The beads were washed once with Wash Buffer and centrifuged at 5,000 x g for 3 min. The beads were resuspended in 2x Laemmli sample buffer and boiled for 2 min to elute bound proteins. Eluted samples and input samples were loaded for western blot analysis.

### Western Blot Analysis

Cells were lysed with RIPA buffer (150 mM NaCl, 10 mM Tris-HCl pH 7.5, 1 mM ethylenediaminetetraacetic acid (EDTA), 1% Triton X-100, 0.1% SDS, 1X supplemented with protease inhibitor and Phosphostop). Then cells were scraped, incubated on ice for 5 min and centrifugation at 16000xg for 10 min at 4°C. Protein concentration was determined using the Precision Red Advanced Protein Assay Reagent (Cat. # ADV02) following the manufacturer’s protocol. 30 μg of proteins were separated by SDS-PAGE using 10 or 14% gel and transferred to nitrocellulose membranes for 120 min at 70V. Membranes were blocked with 5% BSA in Tris-buffered saline +Triton X100 (TBS-T) for 30 min at RT, followed by incubation with primary antibody (see table for antibodies and concentrations) overnight at 4°C. After washing 3x with TBS-T, membranes were incubated for 1hr at RT with Alexa-Fluor conjugated secondary antibodies (see Table). Membranes were then washed again x3 in TBS-T and scanned using the LI-COR Odyssey CLx. All images were analyzed using Image Studio Lite v. 5.2.5.

### Confocal microscopy and image processing

All cell imaging was performed using an inverted Nikon A1R scanning confocal microscope equipped with Perfect Focus and 20x (Numerical aperture NA 0.75) and 40x (oil, NA 1.20) objectives. Image analysis was done with Nikon Elements software and Fiji (version 2.1.0/1.54f). Live cell imaging was performed in a TOCRIS chamber at 37 °C with 5% CO_2_. Imaging was performed at intervals as specified per experiment. Confocal images were collected with z-stacks covering the entire cell height were acquired.26

### Quantification and statistical analysis

Statistical tests performed are described in the figure legends for each graph with comparisons. In general, unpaired t-test, paired t-test, or one-way ANOVA tests were used as appropriate, and performed in GraphPad Prism. Graphs shown also indicate error bar definitions in figure legends (SD, SEM, etc.). p values are listed in figure legends for all comparisons shown on each graph.

## List of inhibitors

List of inhibitors:
| Name | Manufacturer | Concentration | Cat. Number |
| --- | --- | --- | --- |
| Erlotinib | Sigma Aldrich | 10 μM | CDS022564 |
| LY294002 | Cell Signaling | 10 μM | 9901S |
| U0126 | Tocris | 10 μM | 1144 |
| SCH772984 | Selleck Chemicals | 10 μM | S7101 |

### Recombinant DNA

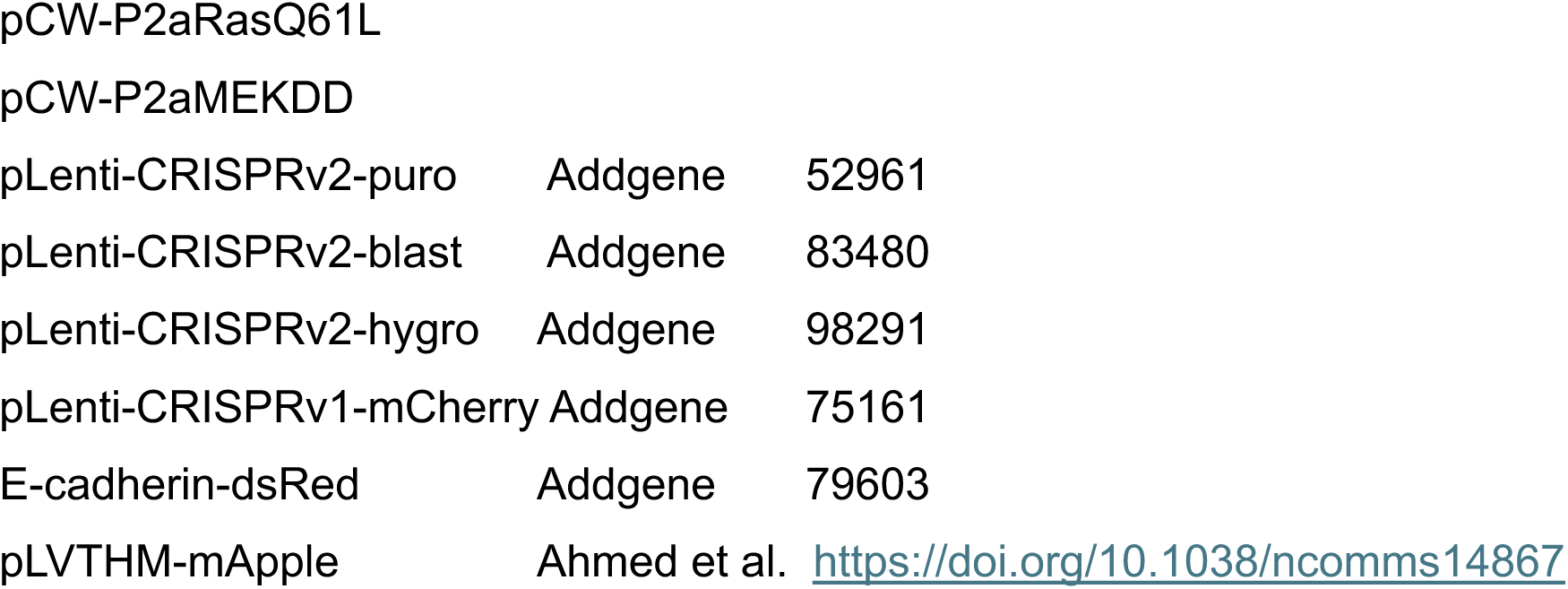

### Oligonucleotides

EGFR_KO_guide CACCGTTCCTCCAACGCCCCACCTG

EGFR_KO_guide CACCGTGAGCCTGTTACTTGTGCCT

SOS1_KO_guide CACCGCTTTTTGTTTACAGGTTCAG

SOS2_KO_guide CACCGTTCTTCGCTGAAGAACTCGT

ECAD_KO_guide CACCGCGTGTCATCAAATGGGGAAG

### Antibodies

Antibodies:
| Name | Manufacturer | Cat. Number | Application | Dilution |
| --- | --- | --- | --- | --- |
| Pan Ras | Calbiochem | OP40 | western | 1:1000 |
| GFP | homemade |  | western | 1:1000 |
| Ran | homemade |  | western | 1:3000 |
| EGFR | EMD Millipore | 06-847 | western | 1:500 |
| PhosphoEGFR | Cell Signaling | 2234S | western | 1:1000 |
| MAPK | Cell Signaling | 4696S | western | 1:1000 |
| Phospho MAPK | Cell Signaling | 4370S | western | 1:1000 |
| AKT | Cell Signaling | 4691 | western | 1:1000 |
| Phospho AKT | Cell Signaling | 4060 | western | 1:1000 |
| SOS1 | Santa Cruz | sc-11793 | Western | 1:500 |
| SOS2 | Santa Cruz | sc-258 | western | 1:500 |
| Tubulin | MilliporeSigma | T9026 | western | 1:5000 |
| GAPDH | Cell Signaling | 2118S | Western | 1:3000 |
| Alexa Fluor 680 donkey anti-mouse IgG (H+L) | Invitrogen | # A10038 | WB | 1:10000 |
| Goat anti-Rabbit (H&L), DyLight 800 4X PEG conjugate | Invitrogen | # SA535571 | WB | 1:10000 |

---

**Figure S1.**
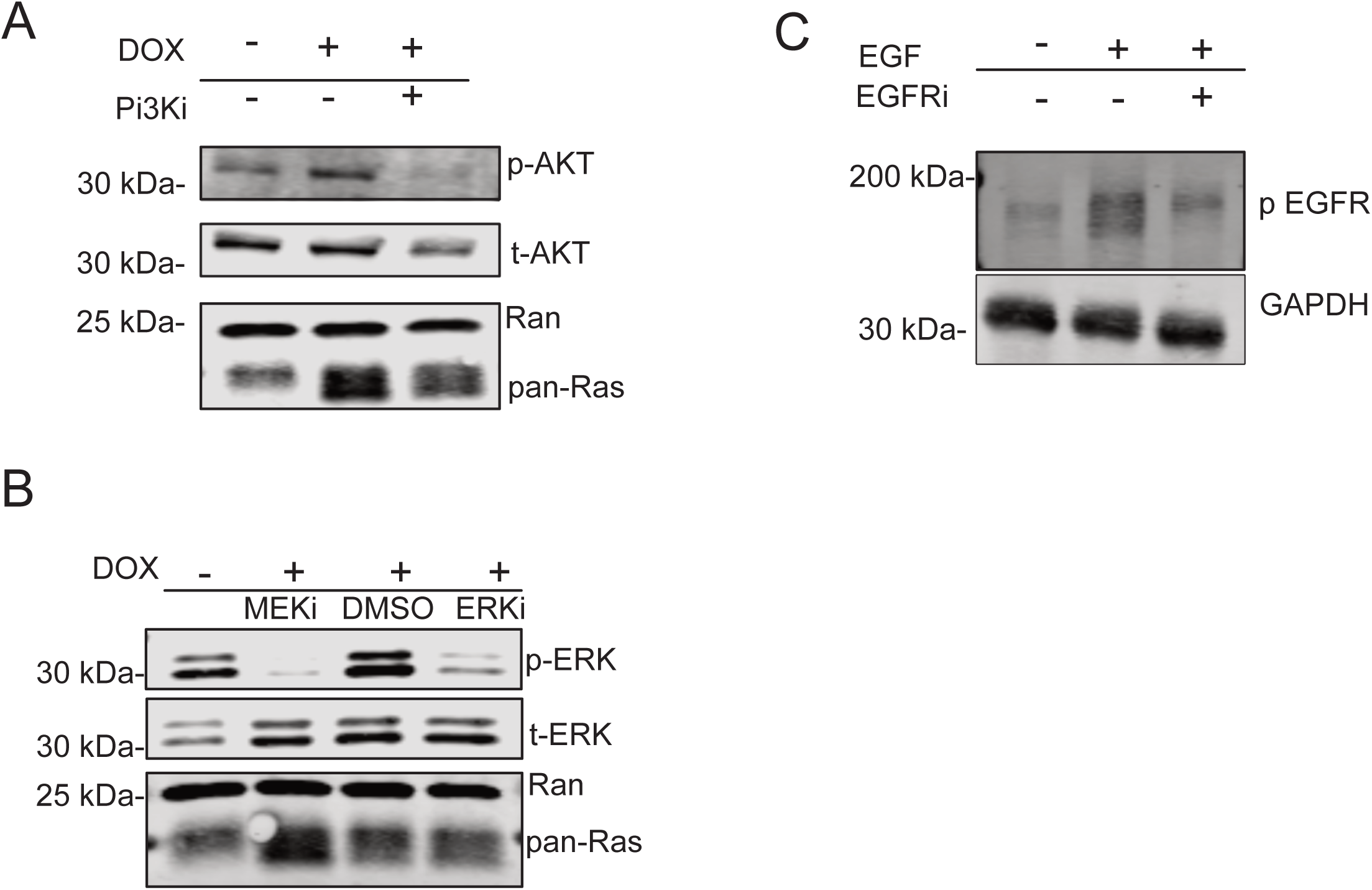
Validation of inhibitors. **(A)**PI3Ki (LY294002) inhibitor suppresses AKT phosphorylation. **(B)** MEKi (U0126) and ERKi (SCH772984) effectively block ERK phosphorylation. **(C)** EGFRi (Erlotinib) reduces EGFR phosphorylation. Representative immunoblots of cell lysates from RasQ61L Eph4 cells +/- Dox.

**Figure S2.**
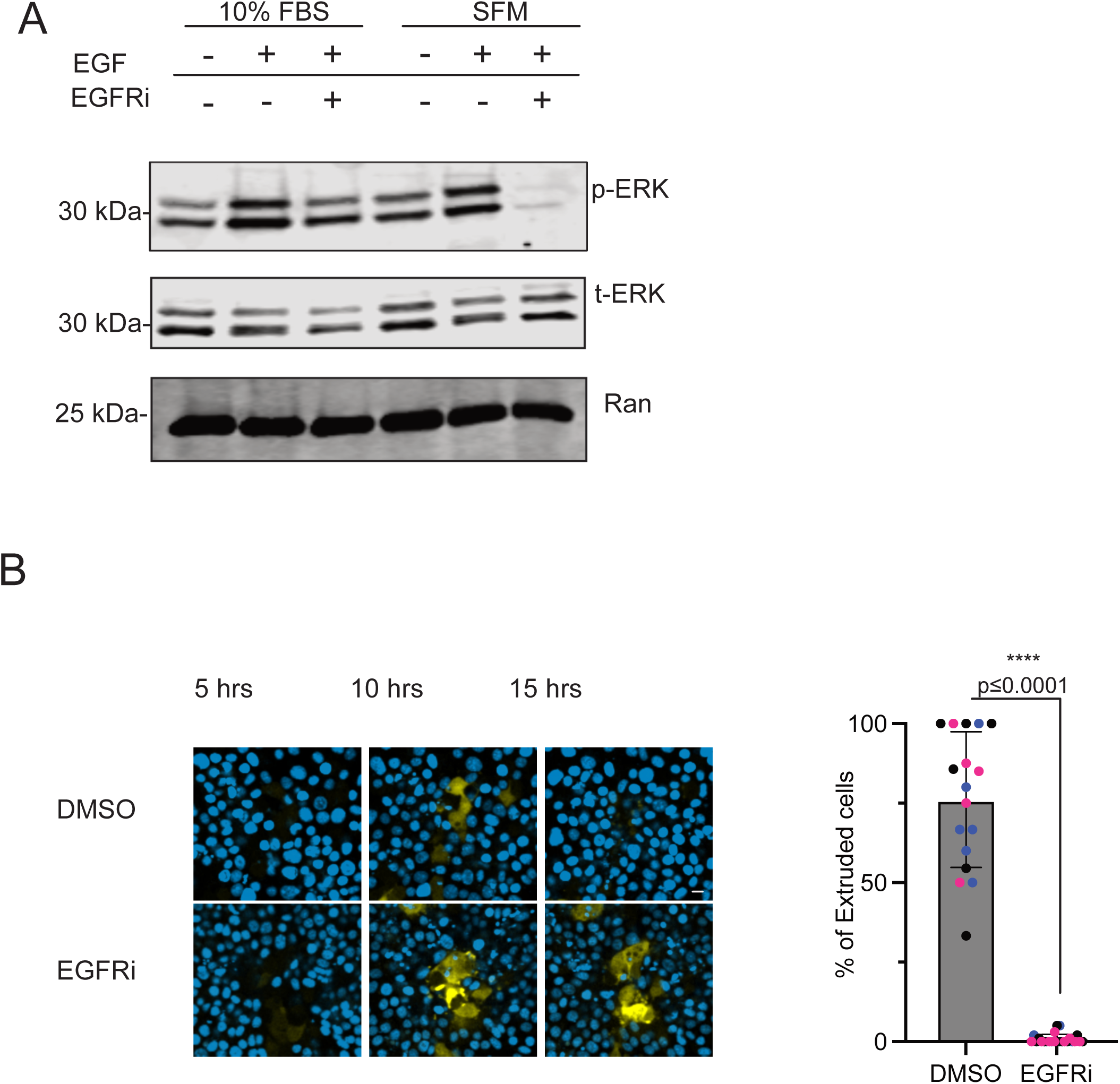
Erlotinib blocks extrusion in the presence of serum, which stimulates ERK phosphorylation independently of EGFR. **(A)**Immunoblot for ERK phosphorylation on lysates of Eph4 cells cultured either in medium with 10% FBS or in serum-free medium (SFM), +/- EGF and +/- Erlotinib to inhibit the EGFR. Note that Erlotinib has very little effect on ERK phosphorylation in FBS, but almost completely blocks ERK phosphorylation in SFM. **(B)** Effect of EGFR inhibition on Ras(Q61L) extrusion in 10% FBS medium. Note that despite the lack of effect on phospho-ERK levels, Erlotinib still blocks extrusion, suggesting that the role of EGFR in extrusion is through a noncanonical pathway independent of ERK phosphorylation. Bars represent mean +/- 1 SD (N=3). Groups were compared using a two-tailed unpaired t-test.

**Figure S3.**
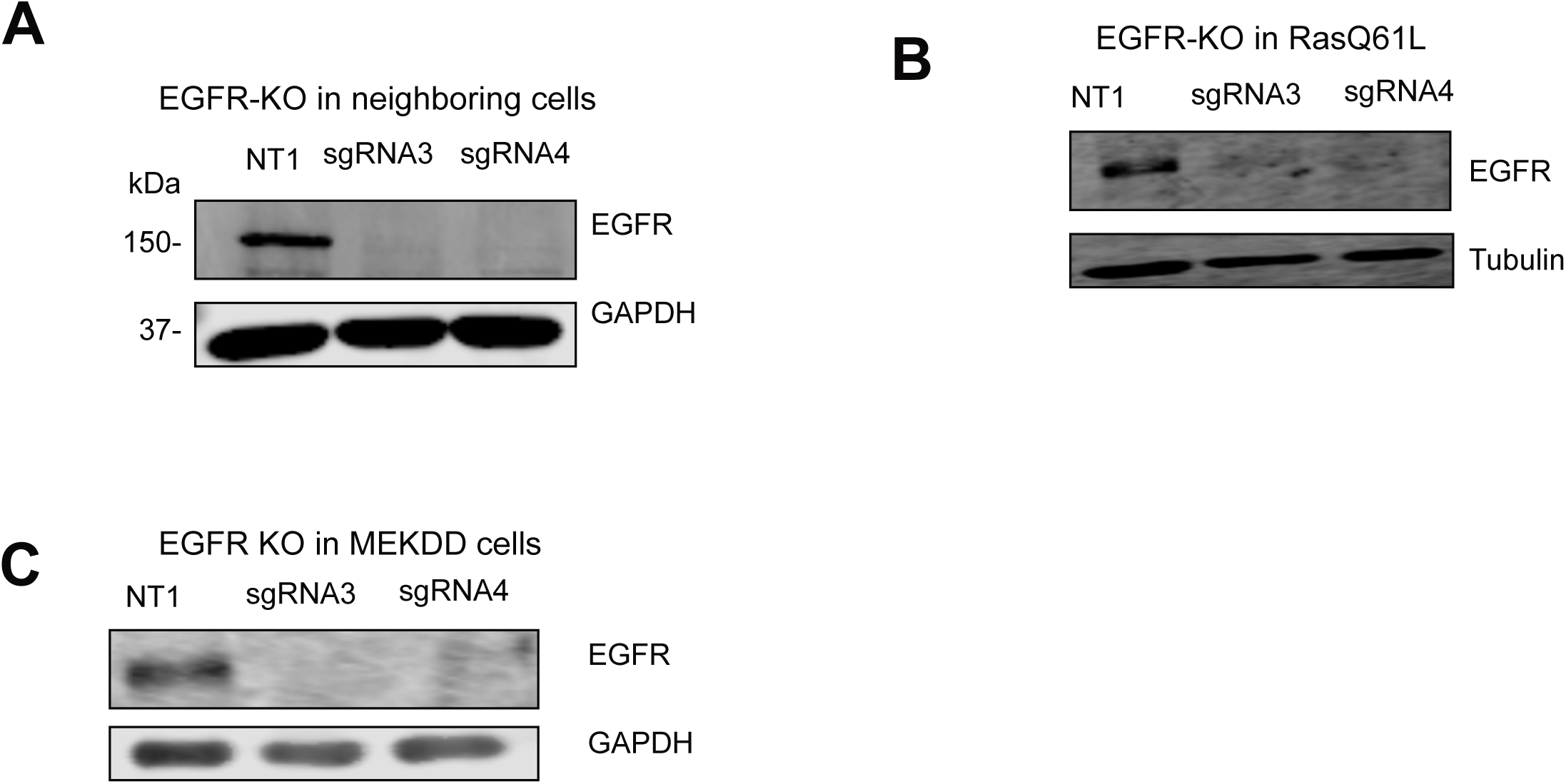
Efficient knockout of EGFR in WT, Ras(Q61L) and MEKDD cells (A-C) Cas9-mediated knockout of EGFR using two distinct sgRNAs. Cell lysates were immunoblotted for EGFR and GAPDH or Tubulin as a loading control.

**Figure S4.**
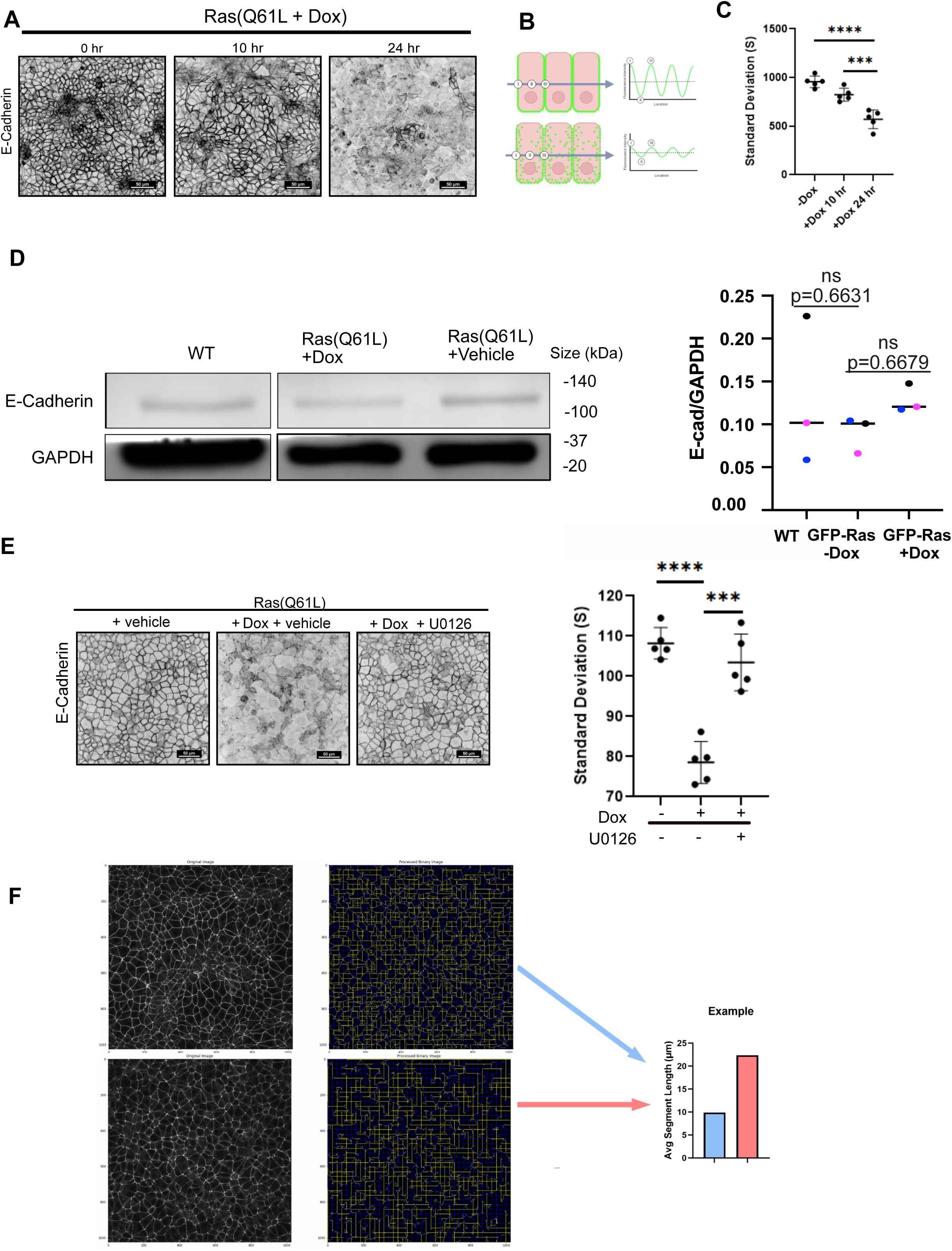
E-cadherin internalization driven by Ras(Q61L) expression. **(A)**Eph4 cells were treated +/- Dox to induce oncogenic Ras and fixed at intervals for staining with anti E-cadherin antibody. Loss of E-cadherin from the adherens junctions is apparent by 24 hrs. **(B)** Lines were drawn across multiple fields of view to measure the variance in staining intensity. Cells with intact junctions show high variance, as compared to those cells in which the E-cadherin has been internalized. **(C)** Quantification is shown of variance across multiple fields of view. **(D)** E-cadherin immunoblot on lysates from WT or Ras(Q61L) cells +/- Dox for 24 hrs. Blots were scanned to quantify band intensities. No significant difference was found between conditions (N=3 biological replicates) showing that the disappearance of E-cadherin from the junctions is not caused by degradation of protein. One-way ANOVA was used to test significance. **(E)** IF images of Ras(Q61L)-expressing cells treated +/- Dox and MEK inhibitor U0126. E-cadherin localization was assessed to evaluate internalization in response to Ras activation. Quantification reflects measurements from multiple independent fields of view. **(F)** Schematic of a robust and quantitative assay for junction integrity, applicable to videos of epithelial cell dynamics. For each video a grid of horizontal and vertical lines is drawn, and for each time point the length of each segment of the grid is determined between pixels of high intensity (top panels white in the figure). To make segment lengths easily visible, they are shown in a different color (yellow or blue) after crossing a junction. As junctions dissolve, the segment lengths increase (bottom panels), as illustrated in the example on the right.

